# Allosteric inhibition of Aurora-A kinase by a synthetic V_NAR_ nanobody

**DOI:** 10.1101/046730

**Authors:** Selena G. Burgess, Arkadiusz Oleksy, Tommaso Cavazza, Mark W. Richards, Isabelle Vernos, David Matthews, Richard Bayliss

## Abstract

The vast majority of clinically-approved protein kinase inhibitors target the ATP binding pocket directly. Consequently, many inhibitors have broad selectivity profiles and most have significant off-target effects. Allosteric inhibitors are generally more selective, but are difficult to identify because allosteric binding sites are often unknown or poorly characterized, and there is no clearly preferred approach to generating hit matter. Aurora-A is activated through binding of TPX2 to an allosteric site on the kinase catalytic domain, and this knowledge could be exploited to generate an inhibitor. However, it is currently unclear how to design such a compound because a small molecule or peptide mimetic of TPX2 would be expected to activate, not inhibit the kinase. Here, we generated an allosteric inhibitor of Aurora-A kinase based on a synthetic, V_NAR_ single domain nanobody scaffold, IgNARV-D01. Biochemical studies and a crystal structure of the Aurora-A/IgNARV-D01 complex show that the nanobody overlaps with the TPX2 binding site. In contrast with the binding of TPX2, which stabilizes an active conformation of the kinase, binding of the nanobody stabilizes an inactive conformation, in which the αC-helix is distorted, the canonical Lys-Glu salt bridge is broken, and the regulatory (R-) spine is disrupted by an additional hydrophobic side chain from the activation loop. These studies illustrate how nanobodies can be used to characterize the regulatory mechanisms of kinases and provide a rational basis for structure-guided design of allosteric Aurora-A kinase inhibitors.

**Significance:** Protein kinases are commonly dysregulated in cancer and inhibitors of protein kinases are key therapeutic drugs. However, this strategy is often undermined by a lack of selectivity since the ATP binding pocket that kinase inhibitors usually target is highly conserved. Inhibitors that target allosteric sites are more selective but more difficult to generate. Here we identify a single domain antibody (nanobody) to target an allosteric pocket on the catalytic domain of Aurora-A kinase and demonstrate that the mechanism is antagonistic to a physiologically-relevant allosteric activator, TPX2. This work will enable the development of allosteric Aurora-A inhibitors as potential therapeutics, and provide a model for the development of tools to investigate allosteric modes of kinase inhibition.

## Introduction

Most current kinase inhibitors target the ATP binding site, which is relatively straightforward to block with small molecules (1). However, this site is also highly conserved among all protein kinases and therefore many inhibitors have off-target effects. Patients treated with kinase inhibitors inevitably relapse due to drug resistance mechanisms such as kinase overexpression, mutation or the activation of bypass pathways, usually involving other kinases. Complex tumour biology, including genetic heterogeneity and drug resistance, may require the inhibition of more than one kinase for effective therapy (2-4). Combinations of kinase inhibitors might address these issues, but combining these drugs safely and effectively is a challenge. This is thought to be, at least to some extent, due to the unfocussed selectivity profiles of ATP-competitive inhibitors. Kinase inhibitors that bind to allosteric sites are more selective than ATP-competitive inhibitors (5). However, allosteric inhibitors are more challenging to develop because kinase targets do not always have a clearly suitable allosteric site, and approaches to targeting such sites though screening and synthetic chemistry are less well developed than for the ATP-binding site. As a consequence, allosteric inhibitors are usually discovered through serendipity and there are few examples of kinase inhibitors of this type in the clinic. More recently, there have been several rational approaches to the development of allosteric inhibitors, which require foreknowledge of suitable binding sites (6-8).

Aurora-A is a Ser/Thr protein kinase that functions primarily in cell division (9). The activity of Aurora-A is stimulated by autophosphorylation of Thr288 in a flexible region termed the activation loop (10). Aurora-A autophosphorylation is inefficient, but is stimulated by TPX2, a microtubule-associated protein that binds to the catalytic domain and stabilises the kinase in an active conformation (11-14). Aurora-A is dysregulated in cancers and has been a popular target for drug discovery (9, 15). The first ATP-competitive inhibitors generated were equally effective against Aurora-B, but there are now a few compounds selective for Aurora-A that have undergone clinical trials such as MLN8054 and MLN8237 (16, 17). To our knowledge, allosteric inhibitors of Aurora-A have not yet been identified and there has been no clearly described plan for their development. The TPX2 binding site is a known site of allosteric regulation, but it is not clear how to design a suitable compound because a small molecule or peptide mimetic of TPX2 would be expected to activate, not inhibit the kinase.

In addition to conventional antibodies, camelids and some sharks have antibodies that consist solely of a heavy chain (18-20). For example, the immunoglobin new antigen receptor (IgNAR) from sharks consist of a heavy chain comprising five constant (C) domains and a single, variable domain, termed V or V_NAR_ that binds targets. V_NAR_ domains have an Ig fold consisting of only 8 β-strands, in which the CDR2 region of a conventional VH domain is replaced by a short β-strand HV2 (21, 22). Camelid heavy chain antibodies also recognise epitopes through a single variable VHH domain. Distinct from the equivalent VH domain of conventional antibodies, the variable domains of heavy chain antibodies are independently stable and retain full affinity for epitope (20). Single domain antibodies, also known as nanobodies, have become popular tools to stabilise proteins and thus facilitate their crystallisation (23-26). Single domain antibodies have also been used as biotechnological tools, forming the basis for sensors that read out the conformational state of the extracellular region of epidermal growth factor receptor (EGFR) (27).

Here, we describe the identification of a V_NAR_ single domain antibody based on a shark heavy chain antibody scaffold that binds and inhibits Aurora-A. The crystal structure of the complex indicates an allosteric mode of action that is antagonistic to the mechanism by which TPX2 activates the kinase. These studies provide a rational basis for structure-guided design of allosteric Aurora-A kinase inhibitors.

## Results

### Identification of a single domain antibody that inhibits Aurora-A

A synthetic library of V_NAR_ domains based on a scaffold isolated from Wobbegong shark was screened for binding to the kinase domain (KD) of human wild-type (WT) Aurora-A. All of the confirmed hits had the same amino acid sequence (Fig. S1A). This protein, which we called IgNARV-D01, was expressed in the periplasm of *E. coli* with a non-cleavable C-terminal His_6_-tag and purified using affinity chromatography and size exclusion chromatography (SEC). IgNARV-D01 was verified to bind Aurora-A by Far Western blotting and SEC (Figure S1B, S2). The affinity of the interaction was determined to be 2 μM by surface plasmon resonance (SPR) (Figure 1A, S3A).

**Figure 1.**
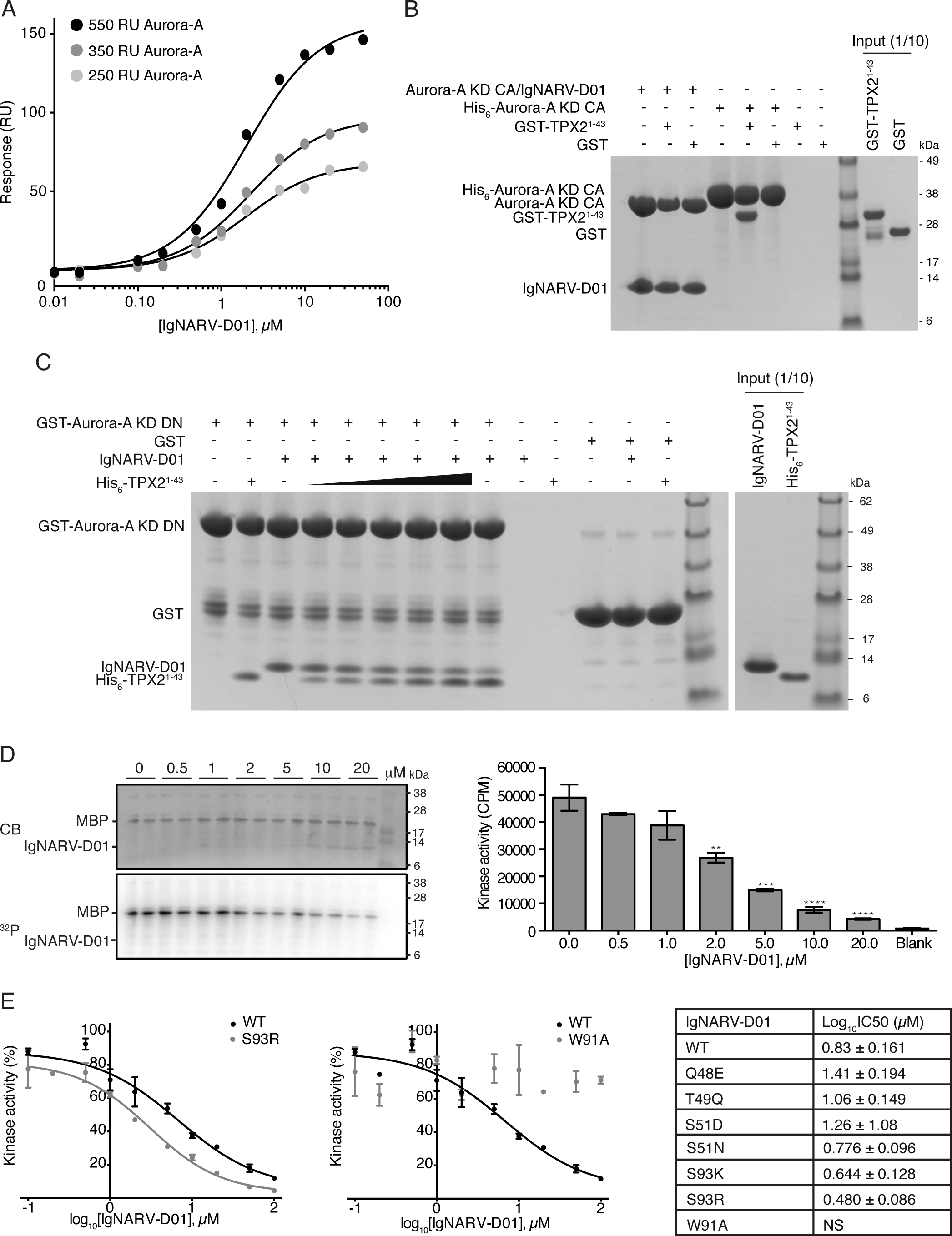
IgNARV-D01 is an Aurora-A inhibitor that competes with TPX2. (A) Surface Plasmon Resonance binding assays between Aurora-A KD CA-Avi and IgNARV-D01. The kinase was immobilised on Biacore Sensor SA chips at 550, 350 and 250 RU and interacted with 0.01 – 50 μM IgNARV-D01. Maximum responses were plotted against IgNARV-D01 concentration and fit to a one-site specific binding equation (black lines) in Prism6 (Graphpad) to calculate binding affinities. (B) Co-precipitation assay between the Aurora-A KD CA/IgNARV-D01 complex or His_6_-Aurora-A KD CA and GST-TPX2 1-43. The complex and Aurora-A were immobilized on Nickel Sepharose beads using the His_6_-tag on the nanobody and kinase, respectively. GST was used as a binding control. (C) Coprecipitation assay between GST-Aurora-A KD DN and IgNARV-D01 and His_6_-TPX2^1-43^. 2 μM GST-Aurora-A KD DN was immobilized on Gluthathione Sepharose 4B beads and incubated with 5 μM IgNARV-D01 and 0, 1, 2, 5, 10, 20 and 50 μM His_6_-TPX2 (black triangle). GST was used as a binding control. (D) *In vitro* kinase activity assay of Aurora-A KD in the presence of IgNARV-D01. MBP was used as a generic kinase substrate. Reactions were analysed by SDS-PAGE (top left panel) and incorporation of radioisotope resolved by autoradiography (bottom left panel). Incorporation of radioisotope was measured by scintillation counting (right). Error bars represent the standard error for two independent reactions. ** = P< 0.01 *** = P<0.001 and **** = P<0.0001 using one-way ANOVA with Dunnett’s post-hoc test compared to the kinase only reaction. (E) *In vitro* kinase activity curves of Aurora-A KD in the presence of WT and mutant IgNARV-D01 proteins. The kinase activity of Aurora-A KD was measured by the incorporation of radioisotope into the generic kinase substrate, MBP by scintillation counting in the presence of 0.1, 0.2, 0.5, 1, 2, 5, 10, 20, 50 and 100 μM IgNARV-D01. Data were normalized to % kinase activity using the Aurora-A KD only reaction as 100 % and plotted against IgNARV-D01 concentration (right). Data were fitted to a log(inhibitor) vs. response – variable slope in Prism6 (Graphpad) to calculate log_10_IC50s (right, solid line). NS = no significant inhibition observed.

We investigated whether IgNARV-D01 affected the interaction of Aurora-A with TPX2 using nickel sepharose to precipitate protein complexes through association with the His_6_ tag attached to a single component (Figure 1B). Here, untagged Aurora-A KD C290A, C393A (CA) was efficiently co-precipitated by IgNARV-D01 but GST-TPX2^1-43^ did not co-precipitate with the complex. In contrast, GST-TPX2^1-43^ was efficiently precipitated by His_6_-tagged Aurora-A KD CA in the absence of the nanobody. This suggested competition between TPX2 and IgNARV-D01 for Aurora-A binding. His_6_-TPX2^1-43^ or IgNARV-D01 robustly co-precipitated with GST-Aurora-A KD D274N (DN) immobilised on glutathione Sepharose beads, but competition was established through a dose-dependent decrease in IgNARV-D01 binding as the concentration of TPX2 was increased (Figure 1C). We used *Xenopus egg* extracts to investigate the competition between IgNARV-D01 and Aurora-A in a situation closer to the physiological pathway (Figure S3B). In this system, TPX2 and Aurora-A robustly interact when extracts are supplemented with RanGTP (11). However, co-precipitation of IgNARV-D01 with *Xenopus* Aurora-A was not observed and we concluded that we would require a nanobody of higher binding affinity and/or generated against the *Xenopus* protein to warrant further investigation.

In light of the competition between IgNARV-D01 and the Aurora-A activator, TPX2^1-43^, we asked whether IgNARV-D01 might also activate the kinase. IgNARV-D01 was added to kinase assays based on incorporation of ^32^P into a substrate protein to quantify the kinase activity of Aurora-A (Figure 1D, 1E). IgNARV-D01 showed a dose-dependent inhibition of Aurora-A, an effect opposite to that of TPX2. To address the mechanism of Aurora-A inhibition by IgNARV-D01, we crystallised the complex in two different forms and determined the structures by X-ray crystallography to limiting resolutions of 1.67 A and 1.79 A, respectively (Table S1). In both crystal forms the asymmetric unit contained a single complex of Aurora-A KD CA/IgNARV-D01 with a 1:1 stoichiometry.

### The binding sites of IgNARV-D01 and TPX2 on Aurora-A overlap

In the crystal structure of the complex, Aurora-A adopts a canonical kinase fold, with a molecule of ADP sandwiched between the N- and C-lobe (Figure 2A). IgNARV-D01 has an Ig fold. IgNARV-D01 makes contacts with both lobes of Aurora-A, but does not closely approach the ATP binding pocket. The interface is centered on the αC-helix within the N-lobe of Aurora-A. This helix within the kinase fold bears residues that are critical for catalysis and is often the site of interactions with regulatory binding partners as exemplified by the complex of Cyclin-A/CDK2, and TPX2/Aurora-A (Figure 2B).

**Figure 2.**
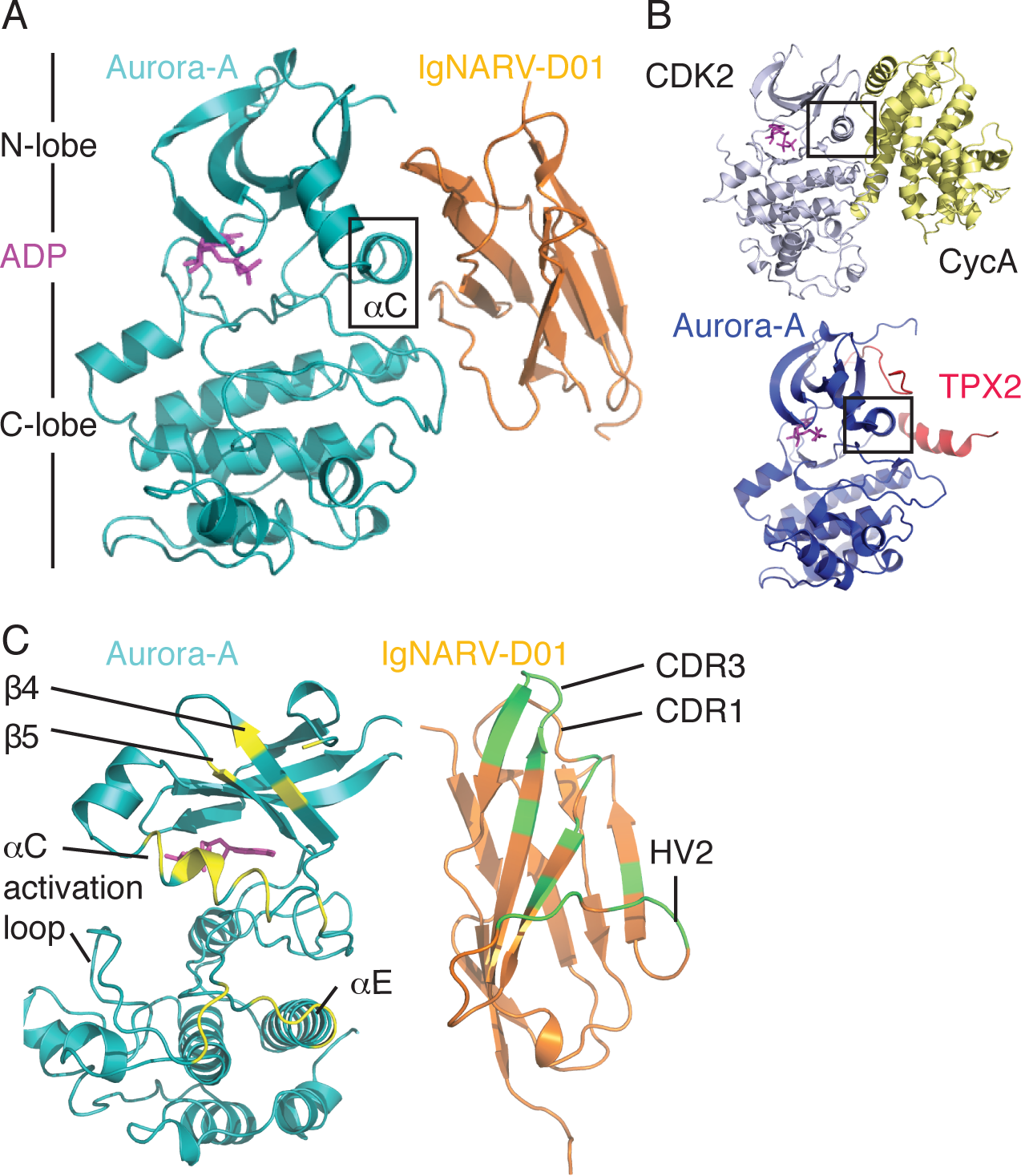
Crystal structure of Aurora-A/IgNARV-D01 complex. (A) Cartoon representation of the complex structure (crystal form 1). Aurora-A is colored teal and IgNARV-D01 is colored orange. (B) Structures of CDK2/Cyclin-A and Aurora-A/TPX2 complexes. The αC-helix is marked with a black rectangle in panels A and B. (C) The interacting regions of the Aurora-A and IgNARV-D01 are shown as contrasting colors on the individual proteins (yellow and green, respectively).

The interface of the complex buries 950 Å^2^ of surface molecule per molecule of IgNARV-D01 or Aurora-A. The binding footprint of IgNARV-D01 on Aurora-A comprises regions of the αC-helix, β4 strand, activation loop and the N-terminus of helix αE (Figure 2C). All three variable regions of IgNARV-D01 contact the kinase surface: Asp33 from CDR1 forms a salt-bridge with Arg179 (Aurora-A αC); residues 48-49 of HV2 contact the N-terminus of αE; Ile52 of HV2 makes side and main-chain interactions with the activation loop sequence Val279-His280-Ala281; the side-chains of CDR3 residues Ile87 and Trp91 insert into a hydrophobic pocket formed between αC and β4, and the Trp91 side-chain makes a H-bond with Glu175 side-chain within this pocket (Figure 3A). In addition, Asn36 and Tyr38 from βC interacts with the αC-helix through a H-bond with the side-chain of Glu183 (and Van der Waals contact with the side-chain of His187). These interactions are mostly conserved between the two crystal forms of the complex, with the exception of the contacts between HV2 and activation loop.

**Figure 3.**
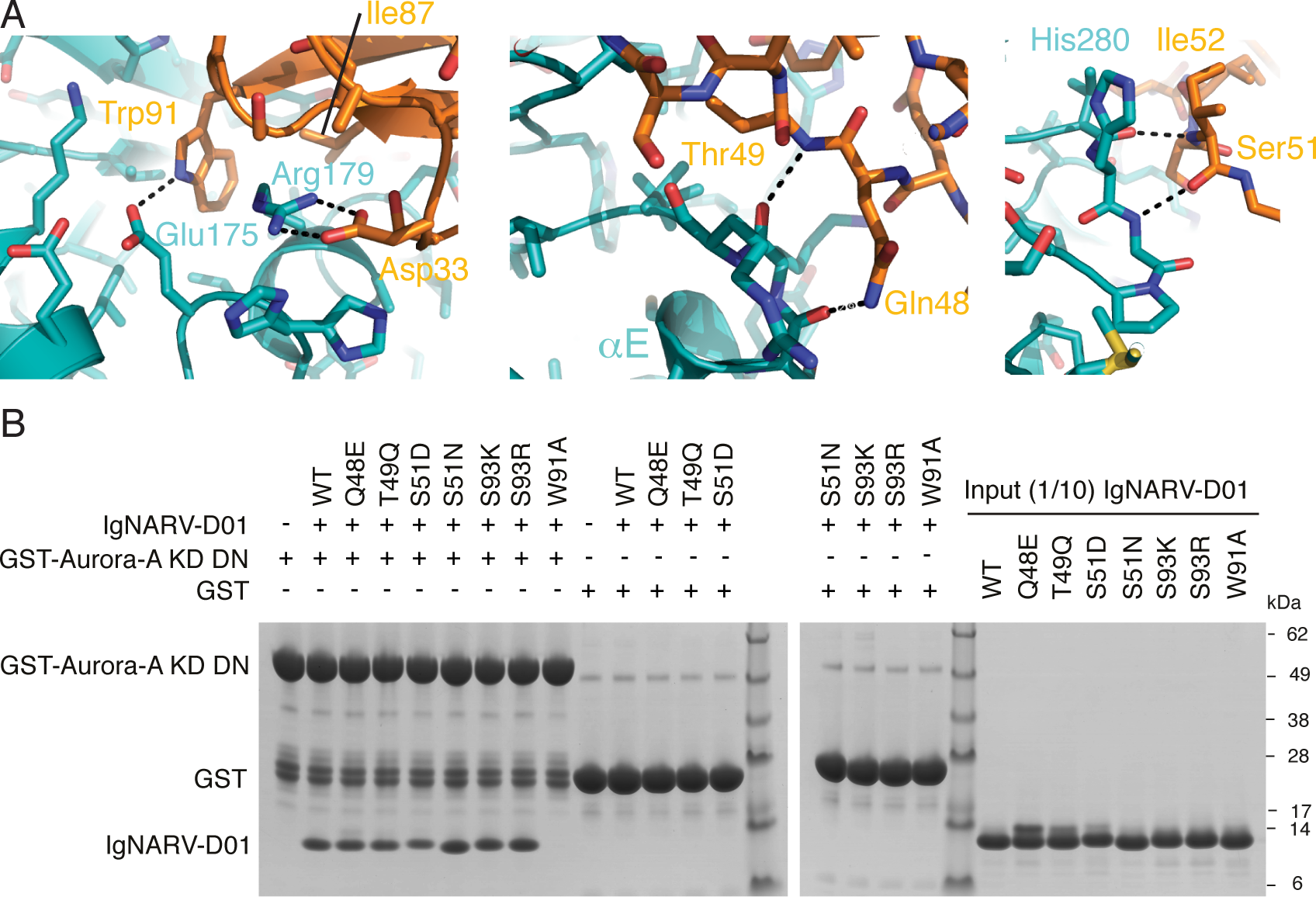
Details of the molecular recognition in the Aurora-A/IgNARV-D01 complex. (A) Key interactions are shown in the three panels. (B) Co-precipitation assay between GST-Aurora-A KD DN and WT, and mutant IgNARV-D01 constructs. GST-Aurora-A KD DN was immobilized on Glutathione Sepharose 4B beads and then incubated with IgNARV-D01 proteins. GST was used as a binding control.

To validate the crystal structure, we generated a point mutation in IgNARV-D01, W91A, designed to disrupt the interaction. We also generated a series of mutations with the aim of enhancing the interaction: Q48E, T49Q, S51D, S51N, S93K, S93R. All mutant IgNARV-D01 proteins were expressed and purified, and tested for binding to Aurora-A by co-precipitation assay (Figure 3B). As predicted, the W91A mutation completely disrupted the interaction and the other mutants retained binding.

We then determined the potency of Aurora-A inhibition by IgNARV-D01 and the point mutants using a kinase assay in which IgNARV-D01 inhibited Aurora-A with an IC_50_ of 6.8 μM (Figure 1E, S4). We could not detect any inhibition of Aurora-A by the W91A mutant IgNARV-D01, consistent with the loss of interaction. Three of the other mutants exhibited less potent inhibition: Q48E, T49Q and S51D; and three of the mutants exhibited more potent inhibition: S51N, S93K and S93R. The most potent inhibitor was S93R-IgNARV-D01, with an IC_50_ of 3.0 μM (Figure S4, S5A).

### IgNARV-D01 destabilises the αC-helix of Aurora-A

The conformation of Aurora-A in complex with IgNARV-D01 lacks key hallmarks of an active kinase: there is no Lys-Glu salt-bridge and the hydrophobic R-spine is incorrectly formed (Figure 4A). In its active conformation, when bound to TPX2, the kinase has a salt-bridge between Lysl62-Glul81, and the R-spine is correctly assembled by the interactions between the side chains of Leul96, Glnl85, Phe275 and His254 (Figure 4B) (14, 28, 29). Indeed, the R-spine of Aurora-A in complex with IgNARV-D01 is disrupted by the presence of an additional side chain: Trp277, which is predicted to interact with substrate peptide in the active kinase, is twisted inwards, and fills the space that Phe275 (of the DFG-motif) would normally occupy. Phe275 is displaced to a position between Lysl62 and Glul81. Trp277 also H-bonds with Gln185, twisting this R-spine component out of position. These conformational changes are coupled to a distortion of the αC-helix in the vicinity of Glu181. The distortion is stabilized by interactions on all sides: the side closest to β4 is stabilized by Trp91 of IgNARV-D01 CDR3; the side facing out to solution is stabilized by Asp33, Asn36 and Tyr38 of IgNARV-D01 CDR1 and strand βC; and the inside is stabilized by conformational changes in the activation loop, most notably Phe275 and Trp277. The position of the αC-helix is incompatible with TPX2 binding, as side-chains of Glul75 and Argl79 would clash with Tyr8 and TyrlO (Figure 4C). In summary, whereas TPX2 stabilises the active conformation of Aurora-A, IgNARV-D01 stabilises an inactive conformation (a schematic overview is shown in Figure 4D).

**Figure 4.**
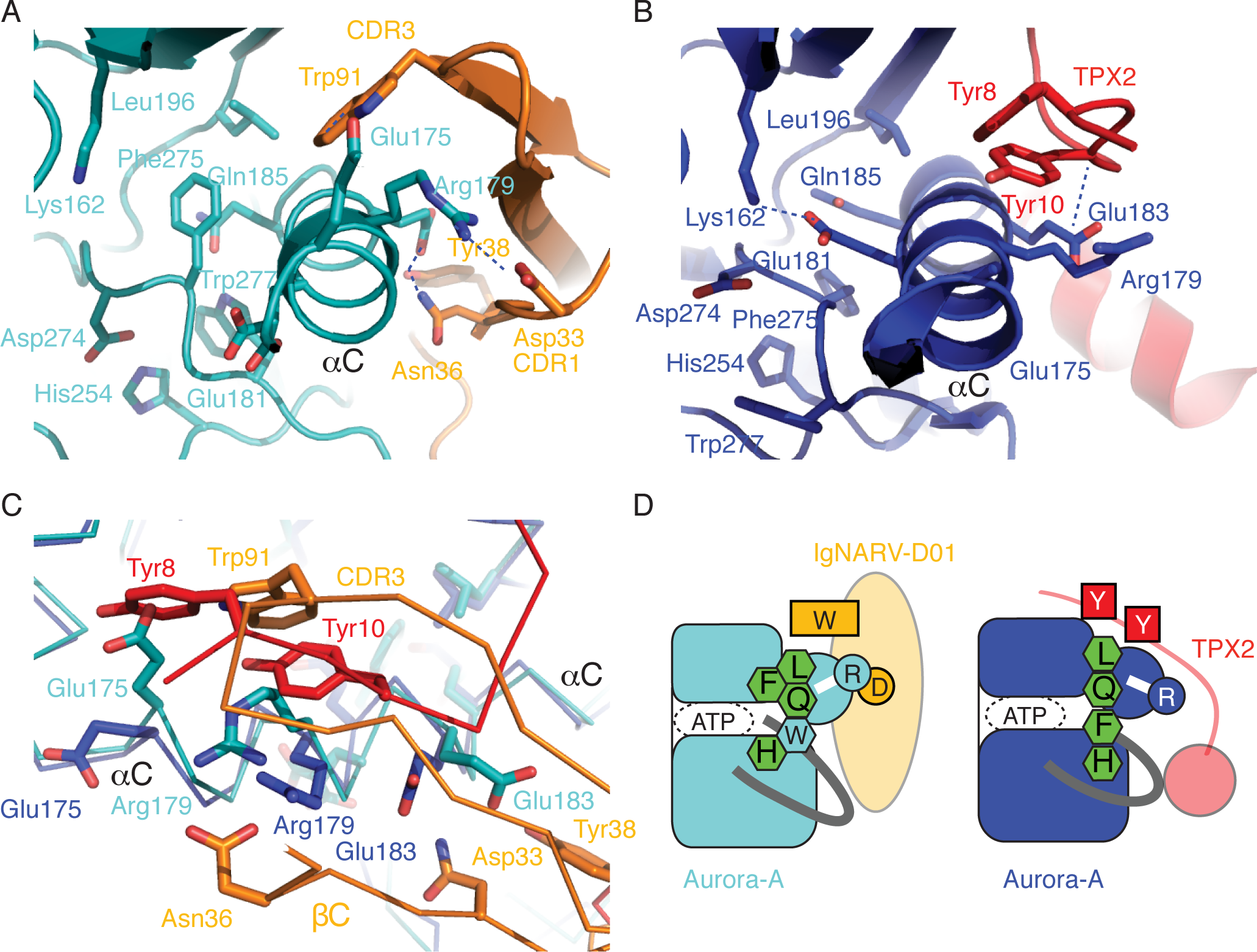
The mechanism of allosteric inhibition of Aurora-A by IgNARV-D01 is antagonistic to TPX2 activation. (A) Aurora-A/IgNARV-D01 complex viewed along the αC-helix. (B) Aurora-A/TPX2 complex, equivalent view to that shown in (A). (C) Superposed structures of Aurora-A/IgNARV-D01 (teal/orange) and Aurora-A/TPX2 (blue/red) complexes viewed with the αC-helix running from left to right Note that the binding site of the CDR3 loop of IgNARV-D01 on Aurora-A overlaps with the binding site of TPX2 residues Tyr8 and TyrlO. (D) Schematic illustration of the structural basis by which IgNARV-D01 and TPX2 stabilize distinct conformations of Aurora-A through binding at the same site. Key residues are shown as single-letter notation. Canonical R-spine residues are shown as green hexagons, and the additional residue that joins the R-spine in inactive Aurora-A is shown as a light blue hexagon.

The conformation of Aurora-A bound to IgNARV-D01 bears a striking resemblance to Aurora-A in complex with MLN8054, an ATP-competitive inhibitor that induces conformational changes in the catalytic domain (Fig. 5A) (16, 30). MLN8054 interacts extensively with the DFG-motif and causes a substantial conformational change in the activation loop. MLN8054 and IgNARV-D01 both stabilise a distortion in the αC-helix that disrupts the Lys-Glu salt-bridge by moving the side chain of Glu181 away from the ATP binding site. The distortion of the αC-helix is coupled to a shift in the position of the side chain of Phe275, which fills the space vacated by Glu181 (Fig. 5B, C). Therefore, the mechanism by which IgNARV-D01 inhibits Aurora-A recapitulates some of the features of MLN8054, without blocking the binding of ATP. Furthermore, the conformation of Aurora-A trapped by IgNARV-D01 is also similar to that observed for Aurora-A in complex with adenosine (Fig. 5D) (31). Thus, three structures of Aurora-A exhibit a distorted αC-helix coupled to changes in the position of Phe275, which represent three distinct complexes of the kinase crystallized under three different conditions. It therefore seems likely the conserved features of these structures represent a physiologically relevant conformation of Aurora-A, one of many that may exist when the kinase is in a dynamic state, which can be captured by ligands or IgNARV-D01.

**Figure 5.**
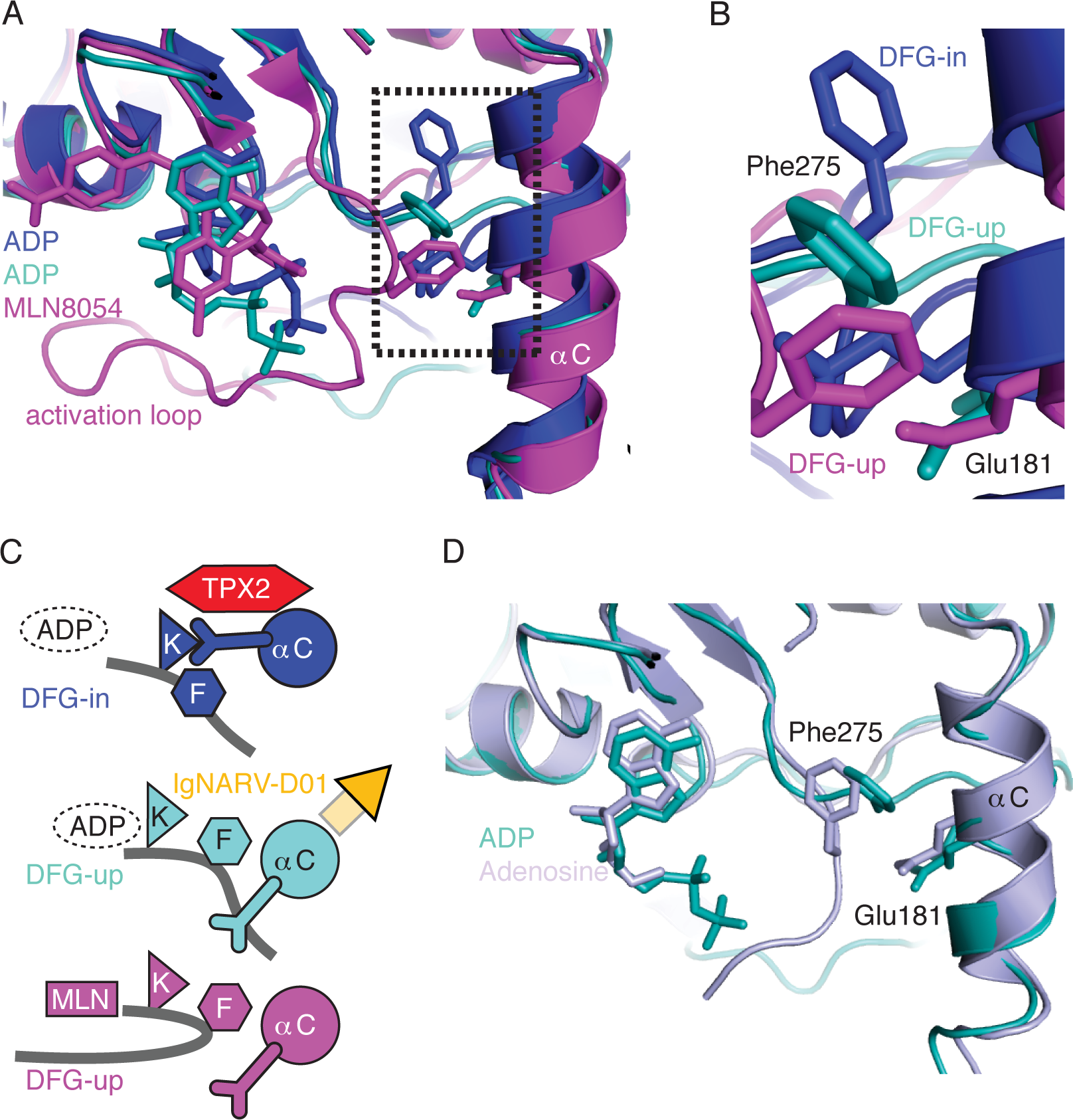
Aurora-A in complex with IgNARV-D01 adopts a DFG-up conformation. (A) Superposed structures of Aurora-A in complex with IgNARV-D01/ADP (teal), TPX2/ADP (dark blue) and MLN8054 (magenta). (B) Magnified view of Phe275 and Glul81. (C) Schematic illustration to show how the DFG-up conformation disrupts the Lys-Glu salt bridge. The activation loop is shown as a grey line, with the position of Phe275 marked with a hexagon labeled ‘F’. In the active conformation of Aurora-A (dark blue, top image), a salt-bridge is formed between Lysl62 (marked with a triangle labeled ‘K’) and Glul81 (shown as a Y-shaped appendage on the αC-helix). Distortion of the αC-helix by IgNARV-D01 (orange, central image) breaks the Lys-Glu salt-bridge and creates a hydrophobic pocket for Phe275. A similar configuration of the αC-helix and Phe275 is observed in the structure of Aurora-A bound to MLN8054 (magenta, lower image), which induces a rearrangement of the activation loop. (D) Superposed structures of Aurora-A in complex with IgNARV-D01/ADP (teal) and adenosine (lilac).

## Discussion

### Single domain antibodies as tools to manipulate kinase structure and activity

Previous work showed that single domain antibodies could be used to trap a kinase in a specific conformation and modulate kinase activity through binding to a regulatory domain (32, 33). Structures of cyclin-G associated kinase (GAK) in two different conformations were captured by co-crystallization with two nanobodies (NbGAK_l and NbGAK_4), which bind to regions of the kinase surface distinct from we observed for IgNARV-D01 (Figure S5B-C) (32). In complex with NbGAK_1, GAK was found in a dimeric arrangement, whereas a dimer was captured in the complex with NbGAK_4. NbGAK_4 modestly enhanced kinase activity, while NbGAK_1 had no effect *Toxoplasma gondii* Calcium-dependent protein kinase 1 (*Tg*CDPKl) consists of a catalytic domain and a regulatory domain that either inhibits or promotes kinase activity in response to calcium. A single domain VHH antibody that interacts with the regulatory domain of *Tg*CDPKl (1B7) inhibits the kinase by stabilising the regulatory domain in the inhibitory conformation (33). These two studies illustrated the potential of nanobodies as tools for biochemical and structural analysis of kinases. Here, we show that the single domain antibody IgNARV-D01 binds directly to the catalytic domain of Aurora-A and inhibits kinase activity through an allosteric mechanism. Taken together, these studies demonstrate the versatility of single domain antibodies as molecular probes to investigate to kinase regulatory mechanisms. However, IgNARV-D01 does not bind Aurora-A with sufficient affinity to be a useful tool in cell-based assays, and we are investigating alternative scaffolds to identify more potent inhibitors.

### Allosteric activators and inhibitors

IgNARV-D01 inserts Trp91 into a hydrophobic pocket formed by the αC-helix and strand β4 of Aurora-A. This pocket plays an important role in the regulation of Aurora-A by TPX2 and, more generally in the regulation of AGC and related kinases, through binding of a peptide bearing a hydrophobic motif (34). These interactions can be in *cis*, as in PKA, or in *trans,* as in Aurora-A, or PDK1, and activate the kinase through stabilisation of an active conformation (35). Specifically, the interaction of TPX2 stabilises the αC-helix of Aurora-A to ensure that the Lys-Glu salt bridge is formed (Fig. 5). This regulatory, hydrophobic pocket presents an attractive target for the rational development of allosteric kinase inhibitors. Indeed, the equivalent hydrophobic (PIF) pocket of PDK1 has successfully been targeted by small molecules, both directly and by tethering approaches (6-8). Here we present an orthogonal approach to targeting this pocket via a single domain antibody. Whereas compounds based on the structure of Aurora-A in complex with TPX2 would be expected to activate the kinase, the Aurora-A/IgNARV-D01 structure could form the basis for rational design of allosteric inhibitors. More generally, we believe that nanobodies will be useful to trap kinases in an inactive conformation to facilitate the development of allosteric kinase inhibitors, and will synergise with other small molecule, peptide and computational approaches.

## Materials and Methods

### Cloning, protein expression and purification

The vectors, pETMll Aurora-A KD, pET30TEV Aurora-A KD CA, pGEX-cs Aurora-A KD, pGEX-cs TPX2^1-43^ and pET30TEV TPX2^1-43^ were produced in earlier work (14, 36, 37). The vector, pGEX-2T was used for the expression of GST. The expression vector for Aurora-A KD-Avi was produced by sub-cloning of the gene for Aurora-A KD into pETM6T1 for expression with a N-terminal TEV cleavable His-NusA tag. A C-terminal non-cleavable Avi-tag was added to the coding sequencing of Aurora-A KD by primer extension PCR.

GST, TPX2^1-43^ and Aurora-A KD, KD CA and KD DN were expressed and purified as stated in earlier work (14, 37). The expression vector for Aurora-A KD-Avi was co-transformed into *E. coli* B834 cells with the vector, pBirAcm for coexpression with biotin ligase and cultured as recommended by the supplier (Avidity LLC, US). His-NusA Aurora-A KD-Avi was purified by immobilized metal ion affinity chromatography (IMAC) using a HiTrap Chelating Sepharose HP column (GE Healthcare) as per the manufacturer’s instructions. The His-NusA tag was removed by overnight TEV cleavage. IMAC was repeated to remove the TEV protease, expression tag and biotin ligase. Q-Sepharose chromatography (GE Healthcare) was performed according to the manufacturer’s instructions to improve protein purity. As a final polishing step, Aurora-A KD-Avi was subject to SEC on a HiLoad 16/600 Superdex 200 column (GE Healthcare) equilibrated in 20 mM Tris pH 7.0, 200 mM NaCl, 5 mM MgCl_2_, 5 mM β-mercaptoethanol and 10% glycerol. Biotinylation of purified Aurora-A KD-Avi was confirmed by Western blotting with an anti-biotin primary antibody (abcam, 1:5000, ab53494).

Expression vectors for IgNARV constructs were transformed into CodonPlus RIL *E. coli* cells and grown in LB media at 37 °C until an induction OD_600_~0.6 was attained and 0.6 mM IPTG added. Cultures were incubated overnight at 21 °C prior to cell harvesting by centrifugation. Protein purification was performed as described in other work for His-tagged constructs (38). The protein was subject to a final SEC step as described for Aurora-A KD-Avi.

### Crystal Structure Determination

To make the Aurora-A KD CA/IgNARV-D01 complex, the proteins were mixed at a stoichiometry of 1:1.2, respectively and were subject to SEC. Fractions containing complex were combined and concentrated to 16.5 mg/ml and incubated with 5 mM ADP/ MgCl_2_ for 1 hour prior to crystallization screening trials. Screens were set-up in 96-well sitting drop MRC plates using a mosquito LCP crystallization robot (ttplabtech) and incubated at 295 K. Crystals were observed after 2 days incubation in a number of conditions. Further hits were identified in the following 3 weeks. Diffraction data was collected at Diamond Light Source (Oxford, UK) on beamline 104-1. Two different space groups were observed and the data from the highest resolution crystals were processed for structure determination. Data used was from a single crystal. Crystal form 1 was produced using 0.2 M ammonium sulfate, 0.1 M sodium acetate pH 4.6, 12.5 % PEG 4000 as the mother liquor. Crystal form 2 was grown using 0.2 M ammonium sulfate, 0.1 M bis-Tris pH 5.5, 25 % PEG 3350 as a precipitant. Both crystals were cryoprotected by the addition of 30% ethylene glycol and flash-cooled. Data processing was carried out using the ‘-3daii’ mode on the *xia2* automated data reduction platform available at Diamond Light Source. The structure of Aurora-A KD CA/IgNARV-D01 was solved by molecular replacement using the structure of Aurora-A KD CA (PDB entry 4CEG) (39) and the structure of a Spotted Wobbegong shark nanobody (PDB 2COQ) (40) as a model. The structure was solved and rigid body refined with PHENIX. Model building carried out with Coot. MolProbity was used for Ramachandran analysis.

Crystals of Aurora-A KD CA/IgNARV-D01 S93R were produced as for the wildtype complex using 0.1 M citric acid pH 5.0, 20% PEG 6000 as a precipitant and diffraction data was collected at Diamond Light Source (Oxford, UK) on beamline 104-1. Structure determination was carried out as described above with data from a single crystal cryoprotected in 30% ethylene glycol using the wild-type complex crystal form 2 as a model.

### Co-precipitation assays

For co-precipitation assays, 100 μg bait protein was immobilized on 20 μl resin equilibrated in assay buffer. The resin was pelleted by centrifugation and washed twice with assay buffer. The beads were resuspended in assay buffer to which 100 μg prey protein was added and incubated for a further 2 hrs at 4 °C. The reactions were washed twice with assay buffer prior to the addition of 20 μl SDS-loading buffer and SDS-PAGE analysis. Gluthatione Sepharose 4B beads (GE Healthcare) equilibrated in the assay buffer, 50 mM Tris pH 7.5, 150 mM NaCl, 5 mM MgCl_2_, 5 mM β-mercaptoethanol and 0.02% TWEEN 20 were used in assays where GST-Aurora-A KD DN and GST were used as bait. Nickel Sepharose resin (GE Healthcare) equilibrated in the assay buffer, 50 mM Tris pH 7.5, 150 mM NaCl, 40 mM imidazole, 0.1 % TWEEN-20 was used to immobilise His_6_-Aurora-A KD CA and IgNARV-D01 as bait proteins.

For competition co-precipitation assays performed with a gradient of 0-50 μM His_6_-TPX2^1-43^ and 5 μM IgNARV-D01, 2 μM GST-Aurora-A KD DN was immobilized on Gluthathione Sepharose 4B beads and reactions were performed as described above.

### In vitro kinase activity assays

Kinase assays were performed as stated in earlier work (37). To determine the IC_50_ of IgNARV-D01 constructs, kinase reactions were performed in the presence of 0 – 100 μM nanobody and analysed by scintillation counting. Data were normalized to % kinase activity using the Aurora-A KD only reaction as 100% and plotted against IgNARV-D01 concentration. Data were fit to a log(inhibitor) vs. response – variable slope in Prism6 (Graphpad) to calculate log_10_IC50.

### Surface plasmon resonance

Surface plasmon resonance assays were performed on a BIAcore 3000 instrument using running buffer, 10 mM Hepes pH 7.4, 150 mM NaCl, 5 mM MgCl_2_, 10% glycerol, 0.005% TWEEN-20. Aurora-A KD-Avi was immobilized on three flow-cells of BIAcore SA sensor chips (GE Healthcare) at three concentrations (250, 350 and 500 RU) and the fourth was left blank. IgNARV-D01 was diluted into running buffer to a range of concentrations and injected over the chips at 40 μL/min for 375 s. Sensorgrams were recorded for each injection and processed using BIAevaluation 3.0 software (Biacore AB) and the data recorded in the blank flow-cell subtracted from each sensorgram. Equilibrium response at 350 s was plotted against concentration for each quantity of immobilized Aurora-A KD-Avi and fitted by non-linear regression to a binding isotherm using Prism6 (GraphPad).

The authors declare no conflict of interest

## Author contributions

S.G.B., A.O., I.V., D.M. and R.B. designed research. S.G.B., T.C. and M.W.R performed research. S.G.B. and R.B. analyzed data. S.G.B. and R.B. wrote the paper.

## Acknowledgements

This work was supported by Cancer Research UK Programme Grant C24461/A13231 (to R.B.). We thank Diamond beamline 104-1 for assistance with data collection.

